# ZOnally magnified Oblique Multi-slice (ZOOM) for cardiac diffusion weighted imaging in vivo

**DOI:** 10.1101/2025.09.20.677511

**Authors:** Lars Mueller, Sam Coveney, Isaac Watson, André Döring, Irvin Teh, May Lwin, Erica Dall’Armellina, Filip Szczepankiewicz, Derek K Jones, Maryam Afzali, Jürgen E Schneider

**Affiliations:** Biomedical Imaging Science Department, Leeds Institute of Cardiovascular and Metabolic Medicine, University of Leeds, Leeds, United Kingdom; Center for Biomedical Imaging, EPFL CIBM-AIT, Lausanne, Switzerland; Dept of Medical Radiation Physics, Lund University, Lund, Sweden; Cardiff University Brain Research Imaging Centre, School of Psychology, Cardiff University, Cardiff, United Kingdom; Department of Cardiovascular Sciences, University of Leicester, Leicester, United Kingdom

**Keywords:** cardiac diffusion MRI, Zonally-magnified Oblique Multi-slice (ZOOM), diffusion tensor imaging, diffusion weighted imaging

## Abstract

**Purpose:** Cardiac diffusion weighted imaging (cDWI) commonly utilizes 2D-selective RF-pulses (such as ZOOMit) to avoid aliasing artifacts when reducing the field of view (FOV). These RF-pulses, which may require parallel transmit capability, typically take several 10’s of milliseconds, prolonging the echo time (TE) of the imaging sequence. Conversely, slice selection using ZOnally-magnified Oblique Multi-slice (ZOOM) (i.e. tilting the excitation relative to the refocusing RF-pulse) is an alternative for reducing FOV while shortening TE. We hypothesized that ZOOM, with appropriate choices for the RF-pulse tilt angle and thickness, can be reliably used as an alternative to ZOOMit for cardiac Diffusion Tensor Imaging (cDTI).

**Methods:** We scanned phantoms using full FOV and ZOOM with various parameters chosen from a 2-dimensional grid of angles and thicknesses. We identified a subset of 5 ZOOM parameters for in vivo cDTI experiments. We performed cDTI on healthy volunteers on a Siemens Prisma (6 subjects) and a Connectom (6 subjects) MR system using the 5 identified ZOOM settings, as well as ZOOMit (Prisma only) and no ZOOM (Connectom only). We compared mean diffusivity (MD), fractional anisotropy (FA), and secondary eigenvector angle (|E2A|) between scans.

**Results:** Average MD, and standard deviation of MD and FA, were significantly reduced by ZOOM compared to no ZOOM on Connectom. We found no significant differences in any averaged diffusion measures between ZOOMit and ZOOM on Prisma.

**Conclusion:** Combining ZOOM with cardiac diffusion weighted imaging allows for TE to be considerably shortened (from 79ms to 67ms) while reducing FOV to avoid aliasing artifacts.

## 1. INTRODUCTION

Diffusion Weighted Imaging (DWI) offers a unique capability to non-invasively assess the micro-structural properties of tissues by sensitizing the signal to the movement of water molecules within tissue ^1^. In vivo human studies of cardiac DWI (cDWI) have shown the ability to characterize the microstructure of the myocardium. ^2^ In particular, cardiac Diffusion Tensor Imaging (cDTI) has been demonstrated to detect alterations in cardiac diseases. It has demonstrated the ability to detect microstructural changes in myocardial infarction, ^3,4,5,6,7^ hypertrophic cardiomyopathy, ^5,8,9,10,11^ aortic stenosis, ^12^ and amyloidosis. ^13^ More recent exploratory work on cardiac Diffusion Kurtosis Imaging has also taken place. ^14,15^. Single-shot spin-echo (SE) echo planar imaging (EPI) with second-order motion compensated gradient waveforms is one of the methods commonly used for cDWI. ^16,17,18^ However, the long readout duration of EPI results in geometric distortion, mostly due to off-resonance effects, and increases echo time (TE), which reduces the signal to noise ratio (SNR). This can be ameliorated with reduced field of view (FOV) acquisitions. However, reducing the FOV in the phase-encoding direction can result in aliasing artifacts as signals from tissues outside the FOV may fold back into the image. Thus, reducing the acquisition FOV requires to limit the excitation spatially.

Limited excitation can be achieved in different ways. ^19^ Suppression-based approaches, such as Outer-Volume Suppression (OVS) ^20,21^ and B1 Insensitive Train to Obliterate Signal (BISTRO) ^22^, saturate signals outside the target FOV before excitation. Other common options are spatially selective RF-based techniques. ^23^ Such approaches include ZOOMit on the Siemens platform, which is a parallel transmit technology (pTX) and the most widely used method for spin-echo-based cDWI . The 2D RF pulse consists of a series of closely spaced sub-pulses combined together. The RF excitation is applied concurrently with a rapidly oscillating gradient along the phase-encode axis and a slower, blipped gradient along the slice-select axis. The slice thickness can be independently adjusted in both the phase-encoding and slice-selection directions. Aligning the rapid gradient along the phase-encoding axis creates a sharp profile and reduces the likelihood of aliasing. The slow (blipped) gradient operates with a much narrower bandwidth, and positioning it along the slice-select axis results in fat-water chemical shift displacements occurring in this direction. By ensuring that the main lobe of the water signal is on-resonance during the refocusing process, fat ghosting artifacts commonly seen in EPI can be significantly reduced. ^24^ However, the duration of these RF-pulses is in the order of several 10’s of ms and can therefore add significantly to the TE and reduce the signal-to-noise ratio (SNR) (the T2 of myocardial tissue is 46 ms at 3T) ^25^.

Recently, cardiac diffusion tensor imaging (cDTI) was combined with ultra-strong gradients. ^26^ This work demonstrated the potential for higher order motion compensation and / or higher b-values but failed to considerably reduce TE, even for more standard diffusion weightings. This was due to the need to acquire a larger FOV to avoid aliasing, since pTX technology (and therefore ZOOMit) is not available on the Connectom MR scanner. Notably OVS was not good enough to suppress the folding artefacts for smaller FOVs. Additional options for reducing the FOV are Inner-Volume Imaging (IVI) ^27^ and Zonally magnified ^28^ methods. These techniques require at least two RF-pulses whose slice orientations are angulated against each other. IVI utilizes refocusing pulses in a plane perpendicular to the excitation pulse, ensuring that only the spins located at the intersection of the two planes contribute to the resulting signal. However, these modifications introduce challenges for multislice imaging, as the perpendicular pulse influences all other slices, if the excitation pulse is angulated like a saturation pulse, and if the refocusing pulse is angulated as an inversion pulse. In both cases this results in strong T1 weighting and almost complete signal cancellation. ^29^

Zonally Oblique Multi-slice (ZOOM) based approaches work by using lower angulation which results in worse outer volume suppression but has less interaction with the other slices, which allows their use in multislice acquisitions. ^29,30,31,32,33,34,35,36^ This approach was previously used in cDWI, ^18,31,36^ but was not investigated further nor were optimized parameters provided. The main parameters to adjust for ZOOM are the angle and thickness of the tilted RF-pulse. The best choice will depend on imaging slice thickness and slice gap, as well as the size of the FOV and the object of interest. In this paper, we investigate optimum parameter setup for a medium angle tilt ZOOM approach that minimizes the amount of cross talk between slices. The optimization is performed for representative geometric parameters commonly used in cDTI. ^2^ We hypothesized that ZOOM, with appropriate choices for the RF-pulse tilt angle and thickness, can be reliably used as an alternative to ZOOMit for multi-slice Cdti

## 2 METHODS

### 2.1 ZOOM theory

The spin echo sequence scheme is shown in Fig 1 . ZOOM can be achieved by angulating either the excitation (RF90) or refocusing (RF180) RF-pulse . The way the reduced FOV is achieved is mostly the same in both methods, but the effects on neighbouring slices are different (as discussed later in this section). The angulated slice direction of one RF pulse is achieved by using a gradient simultaneously in slice and phase direction.

**FIGURE 1.**
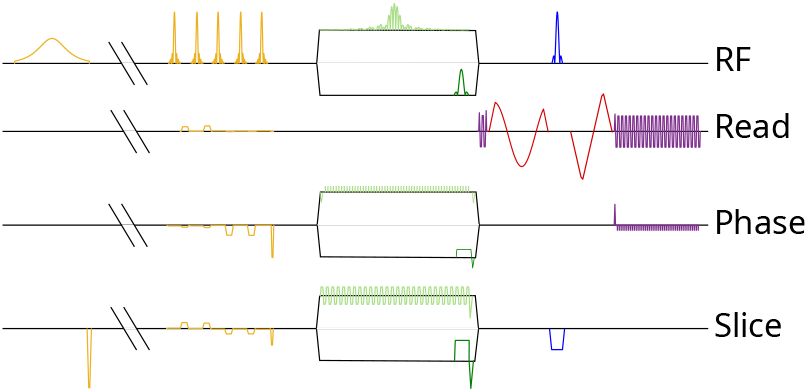
Spin echo sequence scheme for cDTI, including fat saturation and spatial saturation bands (yellow), excitation (light green, top branch: ZOOMit RF pulse with corresponding gradients; dark green, bottom branch: standard RF pulse with slice selection gradients for ZOOM), up to second order motion (i.e. acceleration) compensated diffusion encoding (red, here shown only in read direction), refocusing (blue) and readout (purple). The SPAIR inversion time is cut from the timeline for a more concise presentation.

In Figure 2, three slices are shown, with the acquisition FOV displayed in the center of each slice in lighter (orange) color: the size in phase encode direction is given by FOV_ph_. The angulated slab is described by thickness *τ* and tilt angle *α*. In Figure 2 a, the thickness and angle are such that the entire FOV in the phase encode direction is refocused by the oblique slab. Thus, there is no overlap with the other slices within FOV_ph_, i.e., d_overlap_ = 0; however, d_out_ is large (note that the signal outside FOV_ph_ is not perfectly suppressed by OVS bands, since if it was then there would be no need for ZOOM). A lower *α* than shown in Figure 2 a would only increase d_out_ without giving any advantage for FOV_ph_ (therefore, larger slab thickness increases this minimum angle). For 8 mm slices and slabs, with an 8 mm slice gap, this angle is approximately 4 degrees. In Figure 2 b, the tilt angle is increased such that d_out_ is zero, but this can result in d_overlap_ with the next slices, i.e., cross-talk within FOV_ph_. Only part of the inner volume is fully refocused (FOV_ref_), with partial refocusing occurring in a larger region (FOV_part_). These larger angles are still worth exploring because the real slice profile extends outside the idealized boxes that are shown in Figure 2 .

We therefore have competing goals: to minimize the difference FOV_ph_ – FOV_ref_ ; to minimize d_out_; and to minimize d_overlap_. The best solution is not obvious directly for specific experimental settings with varying OVS effectiveness, real slice profiles and anatomy. Furthermore, there is also the choice for tilting excitation (RF90) or tilting refocusing (RF180), which mostly concerns d_overlap_. For tilting the refocusing pulse the signal in d_overlap_ needs to recover from an inversion until that slice is acquired, whereas for tilting the excitation pulse there is only the need to recover from an excitation into the xy-plane^19^. This will cause a bigger signal loss for RF180 relative to RF90 in most cases.

**FIGURE 2.**
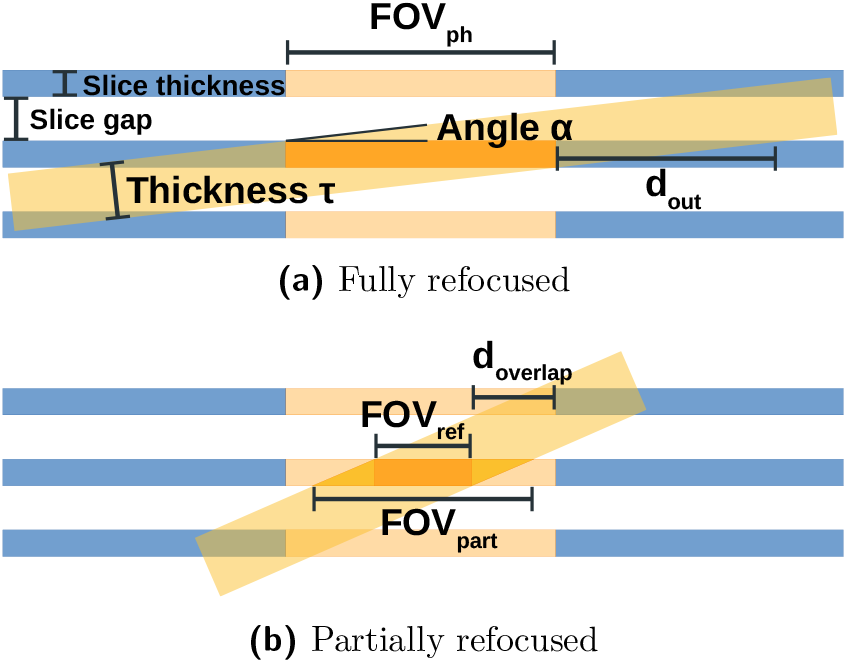
Schematic for ZOOM showing 3 idealized slices. The central light orange region in each slice represents FOV in the phase encoding direction (FOVph). The tilted box represent the oblique slab with thickness but with outer volume the same slice partially excited and angle *a*. (a) Configuration with fully refocused FOV_ph_, but with outer volume in the same slice partially excited d_out_; (b) Inner Volume is fully refocused only in FOV_ref_, and at least partially refocused in FOV_part_. There is no refocusing of the outer volume in the current slice, but there is cross-talk with neighboring slices in d_overlap_.

### 2.2 Parameter optimization

This work investigates ZOOM in the context of cDTI, and therefore the appropriateness of ZOOM parameter choices for slab angle *α* and thickness *τ* are interpreted in the context of diffusion measures obtained from in vivo cDTI using typical acquisition parameters. ^2^ However, it is not feasible to cover the 2-dimensional search space of angle and thickness for a full cDTI in vivo sequence. We therefore opted to select a subset of parameters determined by phantom scans to test in vivo. To simplify the presentation of our work, we provide methodology and results of analyzing the phantom scans as part of the methods, since this informed our choice of parameters for in vivo scans (which constitute our main results). We scanned a spherical water phantom for b-value = 100 s*/*mm^2^ in a single diffusion encoding direction with M2 (including M0 and M1) compensated diffusion gradient waveform, using the geometric parameters as would be used in vivo: 3 slices with 8 mm slice thickness and 8 mm slice gap, with FOV = 320 x 120mm^2^, matrix size= 138x52, and in-plane resolution = 2.32mm^2^. These data were acquired with a prototype sequence on a 3T MRI Prisma and Connectom scanners (Siemens Healthineers, Erlangen, Germany; providing maximum gradient amplitudes of 80 and 300 mT/s and a maximum slew rate of 200 T/m/s) that uses free gradient waveforms (FWF) for diffusion encoding ^37^ and an EPI readout. The sequence further allows to vary the angle and thickness of the slice of the refocusing RF-pulse (Figure 1). The FWF was designed with the NOW toolbox ^38^ up to second-order motion compensation ^39^. Other settings were: TR = 3000 ms, TE = 106 ms (to allow for full field-of-view acquisition), partial Fourier of 7/8, EPI readout, bandwidth per pixel 2130 Hz, with saturation bands along phase encoding direction.

We performed ZOOM with different settings design to cover the 2D parameter space: slab angles *α* (degrees) 0, 4, 5, 10, 15, 20, 25, 30, 90, slab thicknesses *τ* (mm): 8, 12, 16, 20, 24, 28 (i.e. 54 settings in total). We tested tilting excitation (RF90) and refocusing (RF180). We also acquired one full field-of-view data without ZOOM as a reference. Choices of angle *α* are spaced by 5 degrees, but we also include *α* = 4 degrees since this is approximately optimal for *τ* = 8 mm (the smallest slab thickness we test, matching our slice thickness and slice gap). Although angles below *α* = 4 degrees would in theory offer no advantages for *τ ≥* 8 mm, we include *α* = 0 degrees to cover the design space effectively. We defined an ROI inside the phantom of 46 *×* 51 voxels (larger than required to contain almost any left ventricle). The root mean square difference (RMSD) over voxels between the ZOOM image and the reference image was calculated within the ROI as a measure of ZOOM performance.

Figure 3 a shows phantom images acquired using Connectom for a range of combinations of slab thickness and angles, as well as a reference image (zero degree tilt at full FOV). Intensity profiles for the white lines shown in 3 a are shown in Figure 3 b. If the angle is too low (given the thickness), aliasing artifacts are visible in the images and intensity profiles. Figure 4 shows the (log) RMSD results for each parameter combination as colored points, for both Prisma and Connectom. Gaussian process (GP) regression was used to model RMSD as a function of *α* and *τ* (using the software GPy ^40^; radial basis function kernters with automatic relevance determination; hyperparameters were optimized with respect to the log marginal likelihood). The log of the GP posterior mean of RMSD is shown as a heatmap in each subfigure in Figure 4 . We have chosen a color map to help accentuate important features. Notably, the RMSD surfaces all show a ‘valley’ where RMSD is minimized. The position of this valley differs slightly between Connectom and Prisma and between RF 90 and RF 180 but show similar trends. This suggests that we can restrict investigation of parameters in vivo to only one dimension in the parameter space (this is intuitively based on the principles of ZOOM in Figure 2 : larger *τ* requires larger *α*). Furthermore, we decided to limit our in vivo investigation to RF90 only, which showed a smaller RMSD than RF180. We identified 5 co-linear parameters choices for Prisma and Connectom based on this valley in the RMSD surfaces: these choices are shown as white stars in Figure 4 for RF90 subfigures. For Connectom in Fig 4 a: (5deg, 12.5mm), (8deg, 16.4mm), (12deg, 20.3mm), (15deg, 24.1mm), and (18deg, 28mm). For Prisma in Fig 4 b: (3deg, 12.8mm), (6deg, 16.8mm), (8deg, 20.8mm), (11deg, 24mm), and (14deg, 28mm).

**FIGURE 3.**
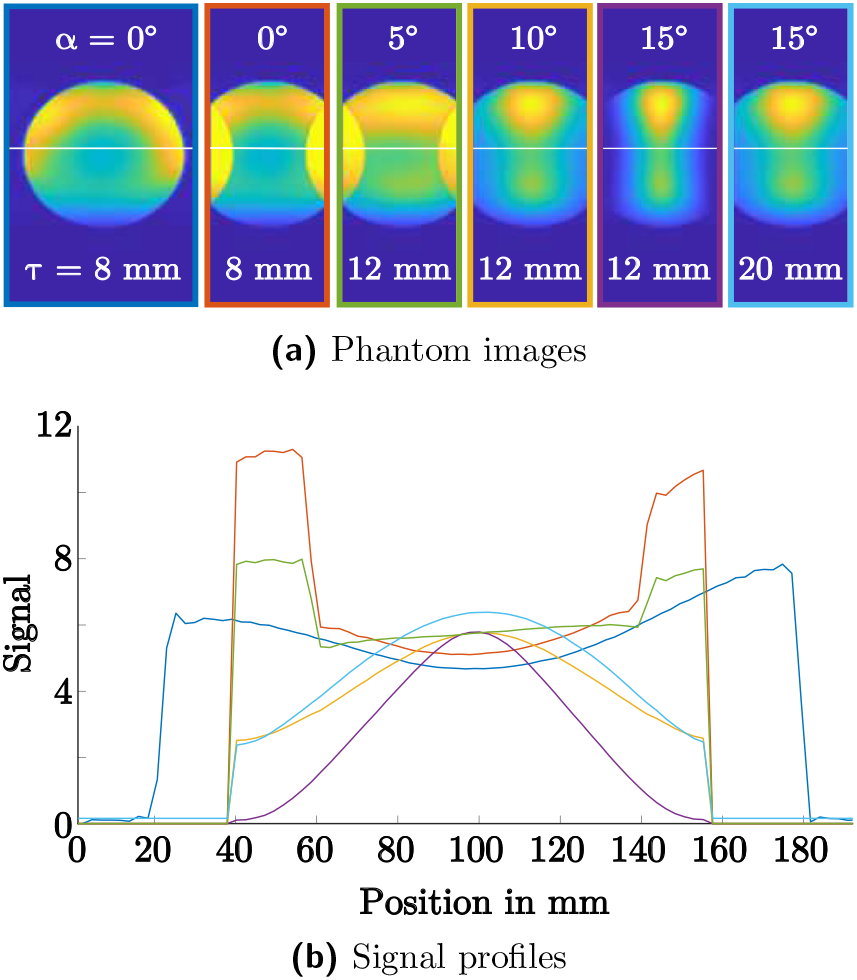
ExamplePimosaigtieosnanind smigmnal profiles from phantom scans on Connectom. (a) Phantom images from Connectom, for a full FOV acquisition (left-most) and several combinations of angle and thickness for the refocusin RF-pulse; (b) Signal profiles along the white horizontal lin shown in (a), drawn with line-colors matching the image borders in (a). Signals were scaled according to the mean over standard deviation in the center of each image.

**FIGURE 4.**
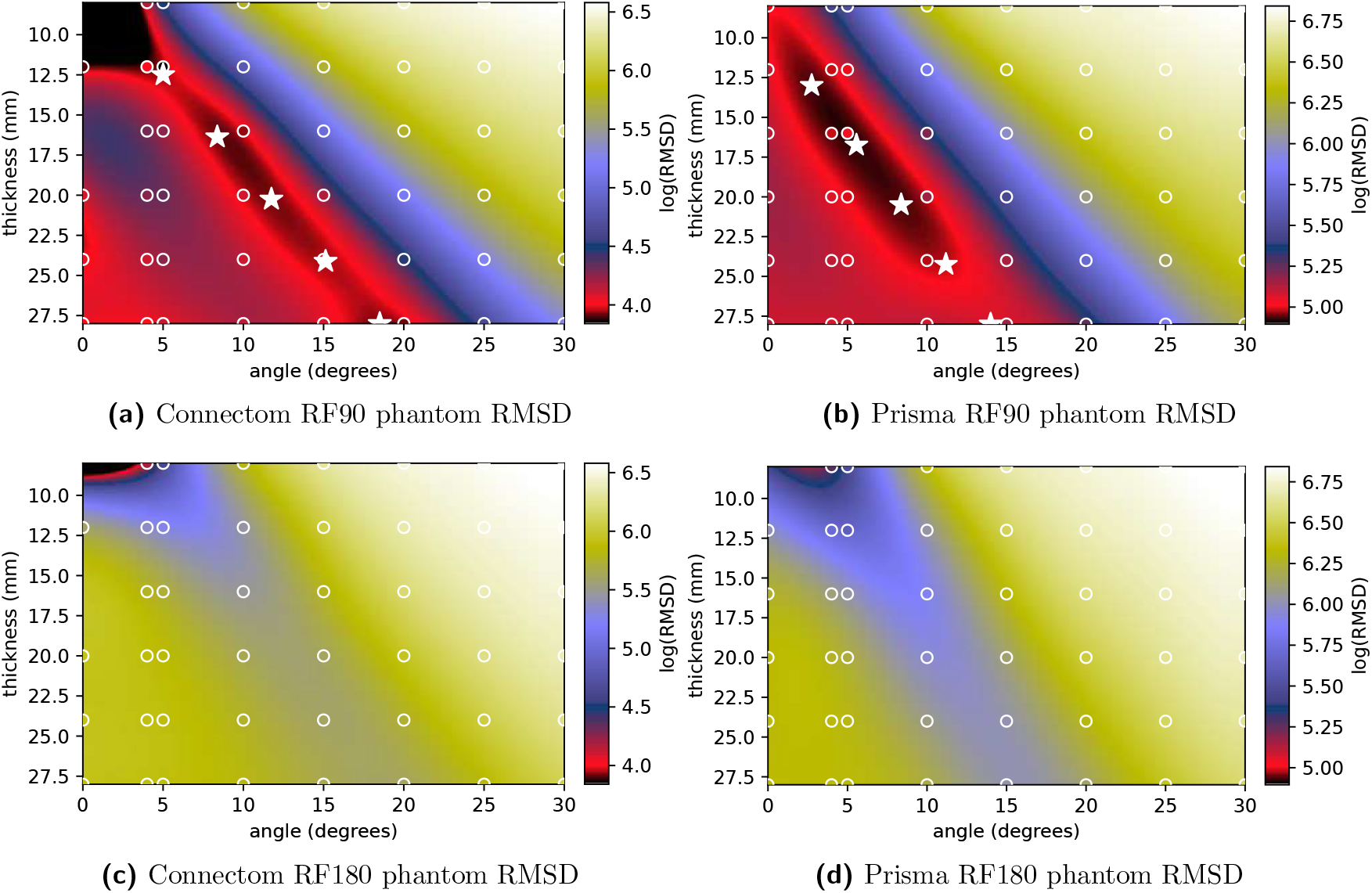
Heatmap of logarithm of *predicted* root mean square difference (RMSD) between a reference scan and ZOOM scans as a function of ZOOM slab thickness and tilt angle, calculated from RMSD ‘observations’ for a subset of angles and thickness (shown as colored points outlined in white) for scans performed on a sphereical water phantom. For RF90, subsets of parameters for testing in vivo are indicated by 5 white stars distributed along the ‘valley’ of minimal RMSD.

### 2.3 In vivo CDTI

Cardiac diffusion-weighted images were acquired for 6 healthy volunteers on Connectom and a different 6 healthy volunteers on Prisma. All volunteers gave written and informed consent. A 18-channel body receive coil and 32-channel spine receive coil were used. The cardiac DWI parameters were TR = 3RR-intervals, TE = 79ms for Prisma and TE = 59ms for Connectom, FOV = 320 x 120mm^2^, matrix size= 138x52, in-plane resolution = 2.32mm^2^, slice thickness = 8mm, partial Fourier 7/8, bandwidth = 2130 Hz/pixel. Note that for Prisma, we matched ZOOM TE to that of ZOOMit to remove confounding factors in our comparison, but it was possible to achieve lower TE of 67ms using ZOOM. Three slices, with 8mm thickness and 8mm gap, were acquired in the short axis plane. Saturation bands were placed around the heart to suppress signal outside the volume of interest. The SPAIR method was used for fat suppression. The diffusion weightings consisted of 3 directions with b = 100 s*/*mm^2^ (12 repetitions) and 30 directions with b = 450 s*/*mm^2^ (6 repetitions). M2 (including M0 and M1) compensated gradient waveforms were used on both MR scanners. ZOOMit was used as the reference scan on Prisma. On Connectom, the reference was without ZOOM. The order of data collection for ZOOM scans was pseudo-randomized across subjects to mitigate the effects of long scan times.

An in-house developed toolbox was used for post-processing. To co-register datasets for each subject together, we first registered a single acquisition: for Prisma this was the ZOOMit scan, and for Connectom this was a chosen ZOOM scan. To register the images in these scans, a user-defined reference image was chosen from the low b-value images and a square mask was defined around the left ventricle. All low b-value images were registered to this reference image, then a new reference image was created from the mean of the registered low b-value images. This new reference image and the square mask was used to register the entire dataset, as well as registering the other acquisitions for the same subject. This guaranteed that all image sets for each subject were co-registered. Registration was rigid and utilized SimpleITK ^41^ with Mutual Information as a metric (calculated within the square mask).

The DTI signal representation was fitted using robust weighted least squares which has been shown to improve fitting results by removing outlier signals. ^42^ The following diffusion measures were calculated: MD, FA, Helix Angle (HA) and secondary eigenvector angle (E2A). HA and E2A utilized a cylindrical coordinate system centered with respect to the left-ventricle. Segmentation was performed on the reference scan that was used to coregister all the datasets for each subject. For each slice, segmentation of the myocardium using manual contours was performed, and regions strongly affected by artifacts (e.g. susceptibility-induced warping) were marked by artifact masks. We then calculated the average and standard deviation of DTI measures over myocardial voxels for each subject and each scan, excluding the artifact regions since they do not represent tissue properties. To determine whether there were any significant differences over subjects between types of scan, average and standard deviation of diffusion measures were compared between scans using paired t-tests. Given the results of these statistical tests, we opted not to apply correction factors for multiple-comparisons, in order to reduce the probability of Type 2 errors (missing a significant difference where one exists), which we feel is justified given the results we obtained.

## 3 RESULTS

Figure 5 a shows example in vivo DWI images for 3 slices on Connectom without ZOOM (reference) and with ZOOM (*α* = 12deg, *τ* = 20mm). There are visible aliasing artifacts in the acquisition without ZOOM that are not present in the ZOOM images. Figure 5 b shows example in vivo DWI images for 3 slices on Prisma with ZOOMit (reference) and with ZOOM (*α* = 12deg, *τ* = 20mm). In the region of interest (i.e. the left ventricle) no aliasing artifacts are visible in either case.

**FIGURE 5.**
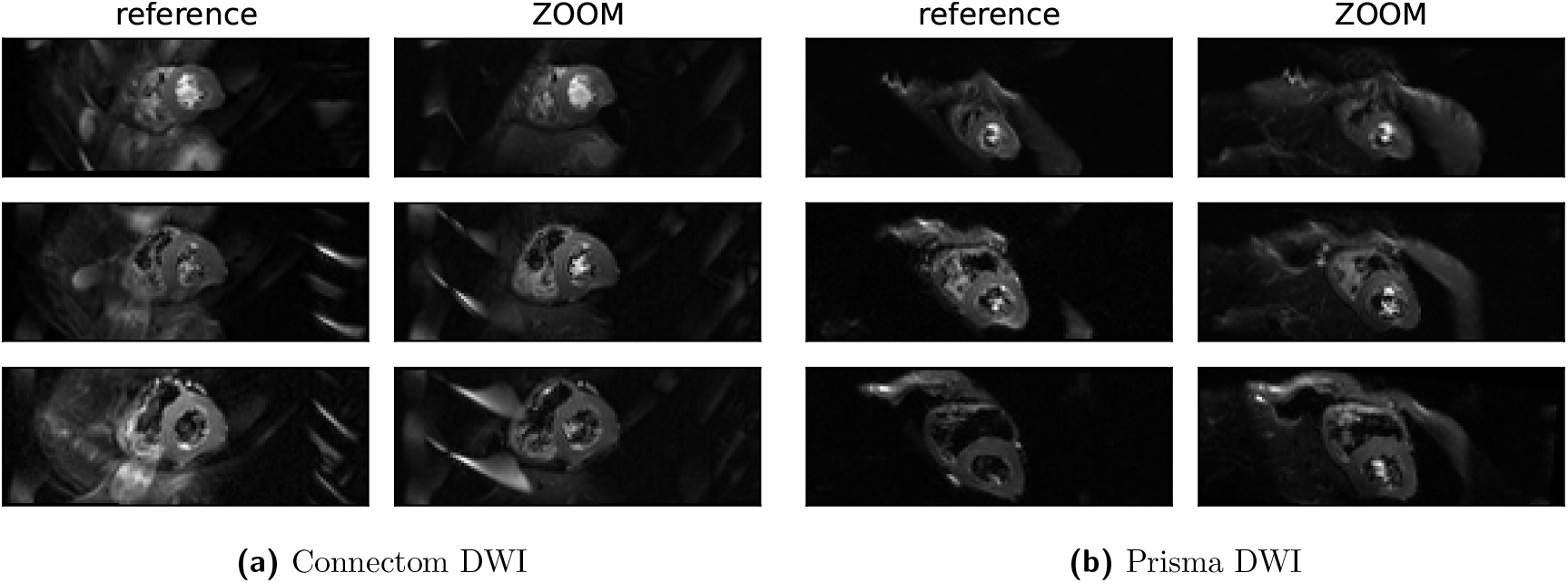
Representative diffusion weighted images acquired on (a) Connectom (reference no ZOOM; ZOOM with *α* = 12*°, τ* = 20mm) and (b) Prisma (reference ZOOMit; ZOOM with *α* = 8*°, τ* = 20.8mm). For each scanner, the reference and ZOOM scans how the same subject (top: apical, middle: mid, bottom: basal). The reference (no ZOOM) scan on Connectom shows aliasing artefacts that are not present with ZOOM.

Figure 6 shows DTI parameter maps for the same subject and acquisitions as Figure 5 a, with the reference (no ZOOM) acquisition in Fig 6 a and the ZOOM acquisition in 6 b. Without ZOOM, the MD values are higher than reported in literature, and highly inhomogeneous. Furthermore, FA is reduced by ZOOM in the same regions where MD is elevated without ZOOM. Both observations suggest artifacts are responsible. The HA transmural pattern is much clearer in 6 b. Note that there are susceptibility artifacts (warping) in both sets of acquisitions. **8** et al Figure 7 shows DTI maps for the same subject and acquisitions as Figure 5 b, with the reference (ZOOMit) acquisition in Fig 7 a and the ZOOM acquisition in 7 b. In this case, the DTI maps look very similar (the HA distribution looks clearer for the apical slice in the ZOOM acquisition than for ZOOMit). As in the case of Fig 6, there are still susceptibility artifacts caused by EPI.

**FIGURE 6.**
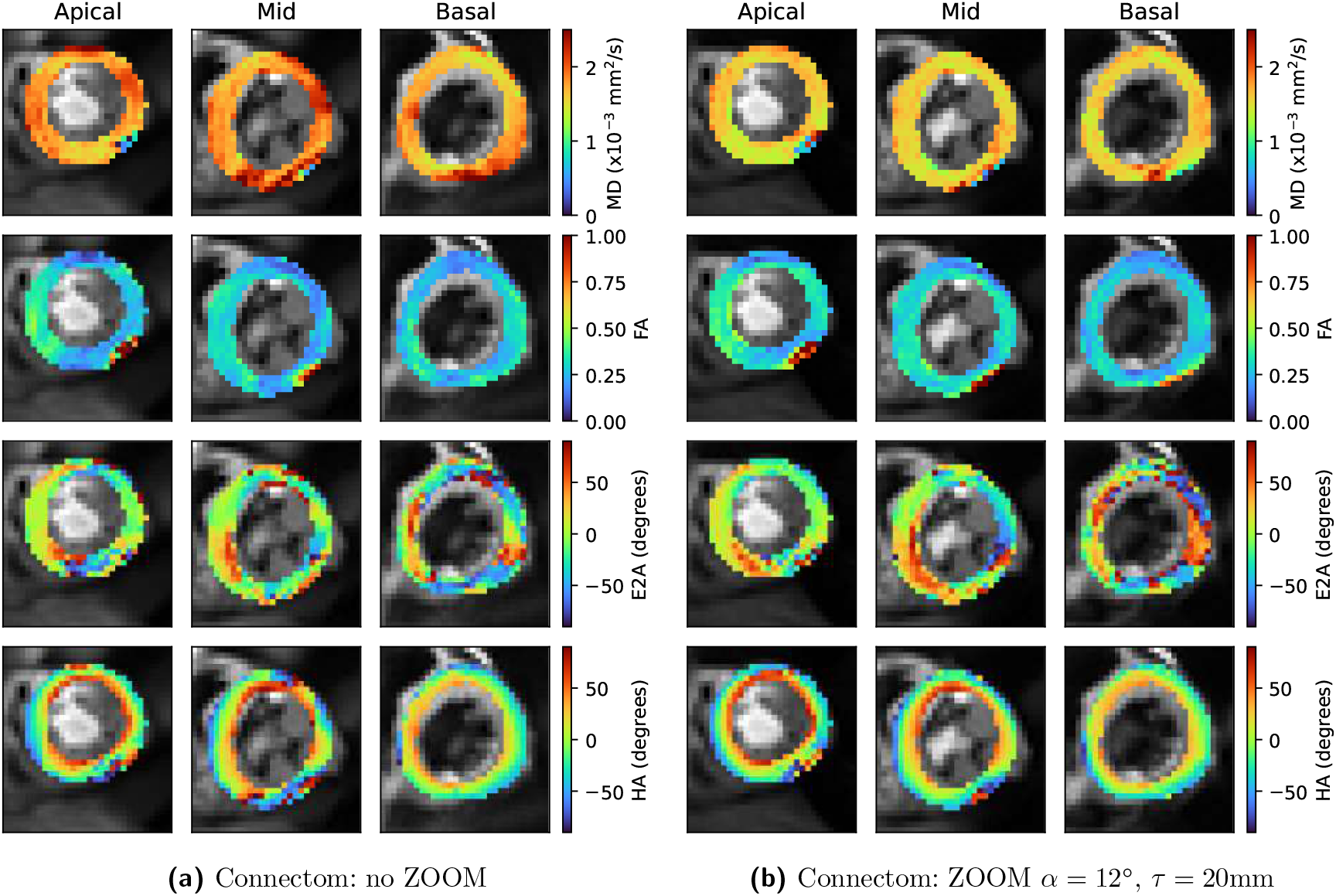
DTI parameter maps acquired for (a) Connectom without ZOOM (b) Connectom with ZOOM. ZOOM reduces MD considerably into a physiologically plausible range, and also increases FA.

**FIGURE 7.**
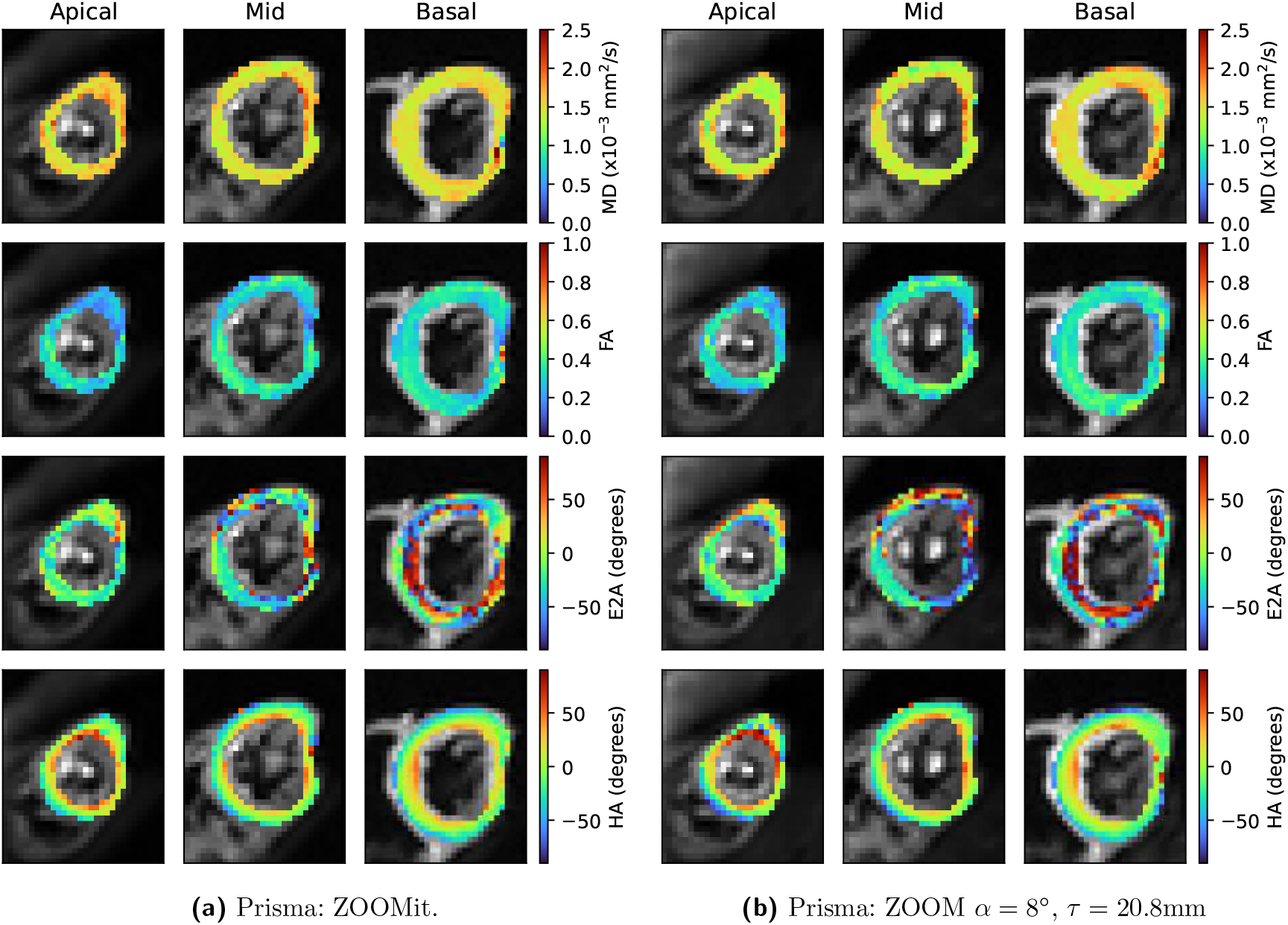
DTI parameter maps acquired for (a) Prisma with ZOOMit (b) Prisma with ZOOM. The parameter maps are very similar in both cases, with no obvious regions of aliasing artefacts.

Figure 8 and 9 show box plots of the average and standard deviation, respectively, of cardiac DTI measures over myocardial voxels over each slice (showing |E2A| rather than E2A) for 6 subjects (i.e. 18 points), for the five different ZOOM schemes tested on each scanner, as well as no ZOOM for Connectom and ZOOMit for Prisma. Figure 10 shows the p-values for paired t-tests on the average and standard deviation of DTI measures between all acquisitions on each scanner. On Connectom, Figure 8 a shows that all ZOOM settings result in average MD values that are reduced compared to the no ZOOM setting, and the first row (or column) of 10 a shows that these results are statistically significant. Furthermore, the spread of measures (over subjects) is less for MD and FA with ZOOM compared to no ZOOM. Figure 10 a shows that the standard deviation over voxels with ZOOM is reduced compared to no ZOOM for both MD and FA, representing a reduction in regions with artifacts in each subject caused by aliasing (as can be seen in the example Fig 6 a).

**FIGURE 8.**
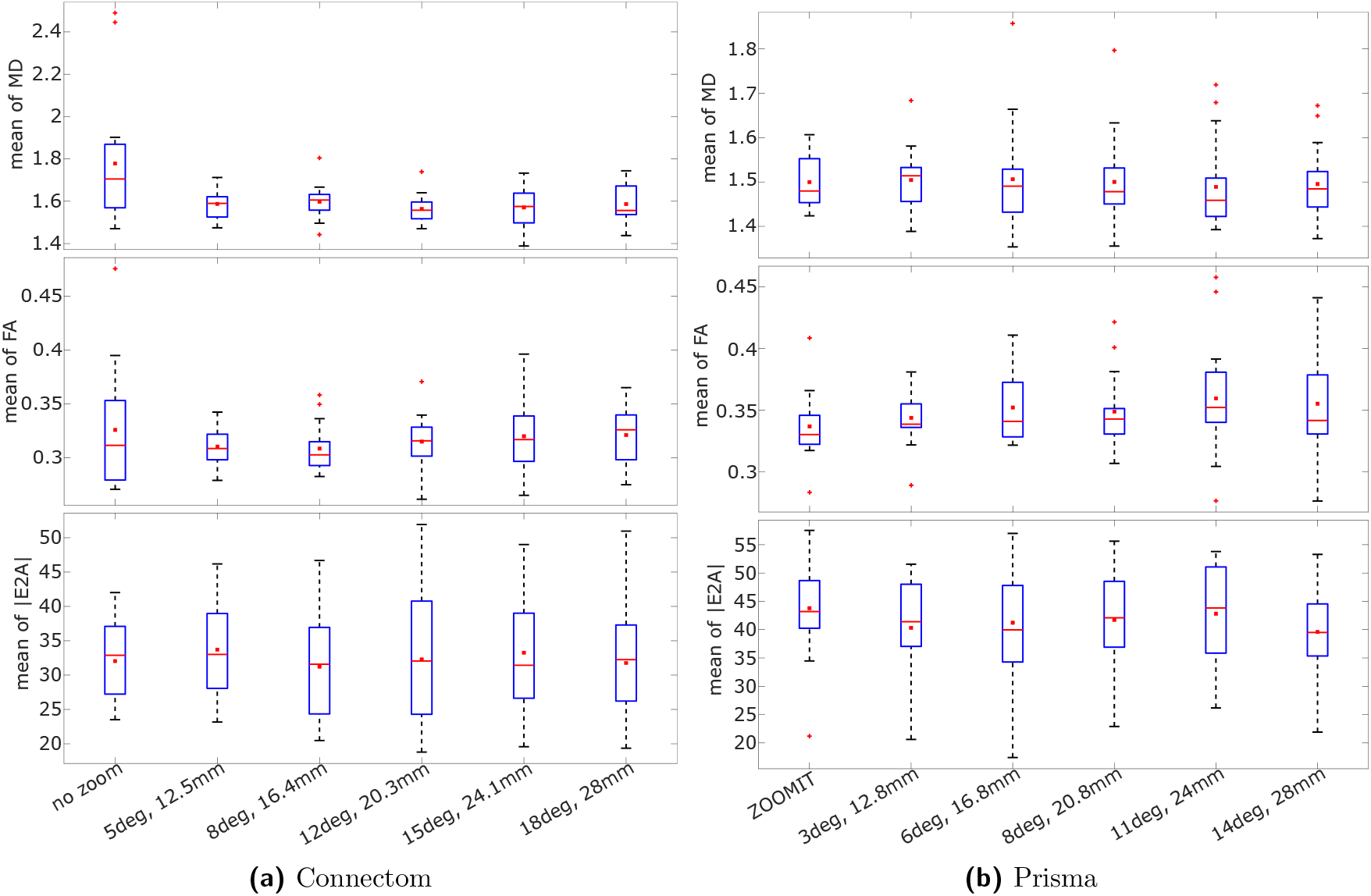
Box plots of average (over myocardial voxels) of mean diffusivity (MD, *×*10^-3^mm^2^/s), fractional anisotropy (FA), and absolute secondary eigenvector angle (|E2A|, degrees), for six subjects for Connectom and Prisma. (a) No Zoom followed by five different ZOOM schemes for Connectom; (b) ZOOMit followed by five different ZOOM schemes for Prisma.

**FIGURE 9.**
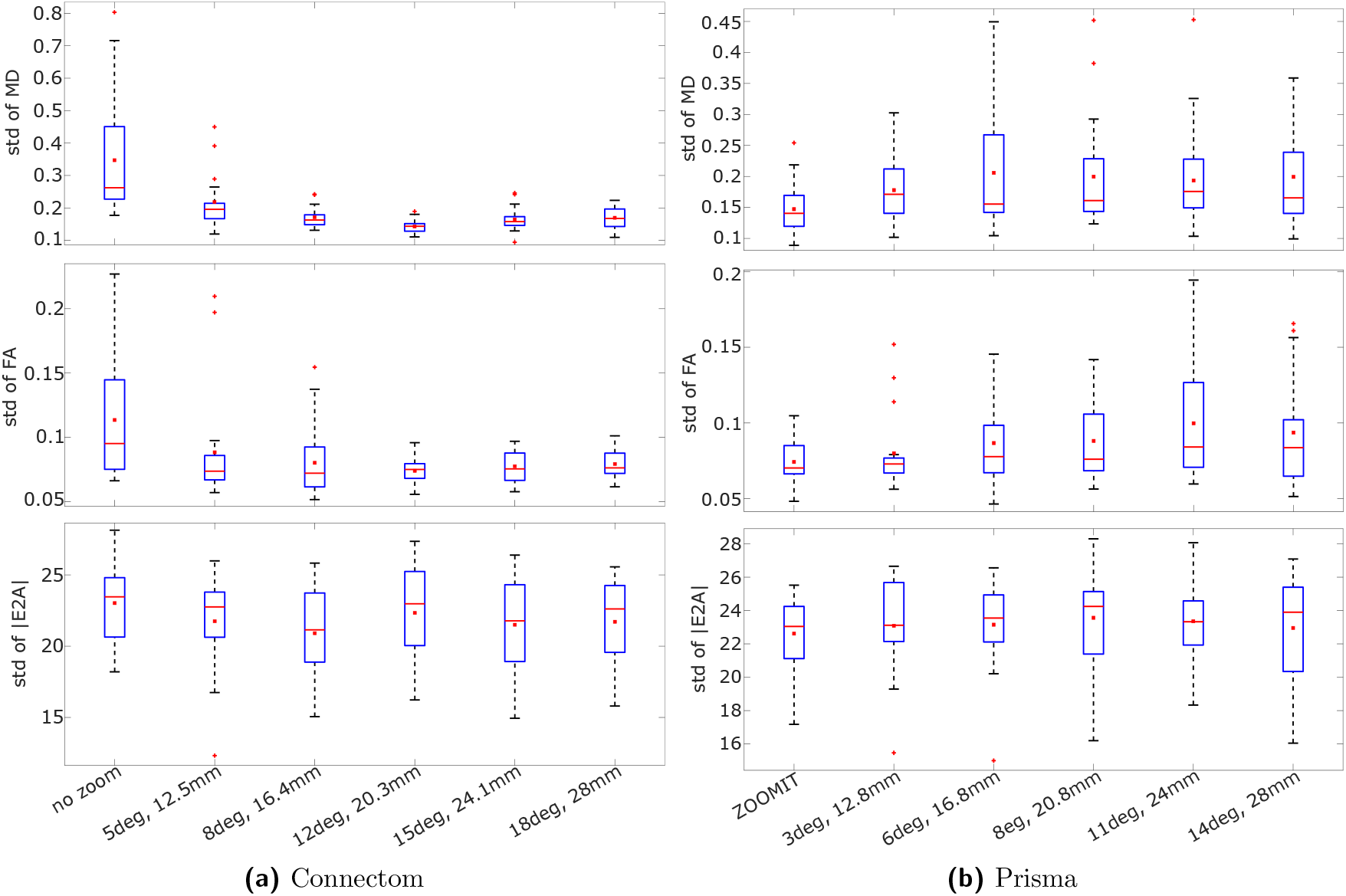
Box plots of standard deviation (over myocardial voxels) of mean diffusivity (MD, *×*10^-3^mm^2^/s), fractional anisotropy (FA), and absolute secondary eigenvector angle (|E2A|, degrees), for six subjects for Connectom and Prisma. (a) Five different ZOOM schemes and one without ZOOM for Connectom; (b) Five different ZOOM schemes and one for ZOOMit for Prisma.

**FIGURE 10.**
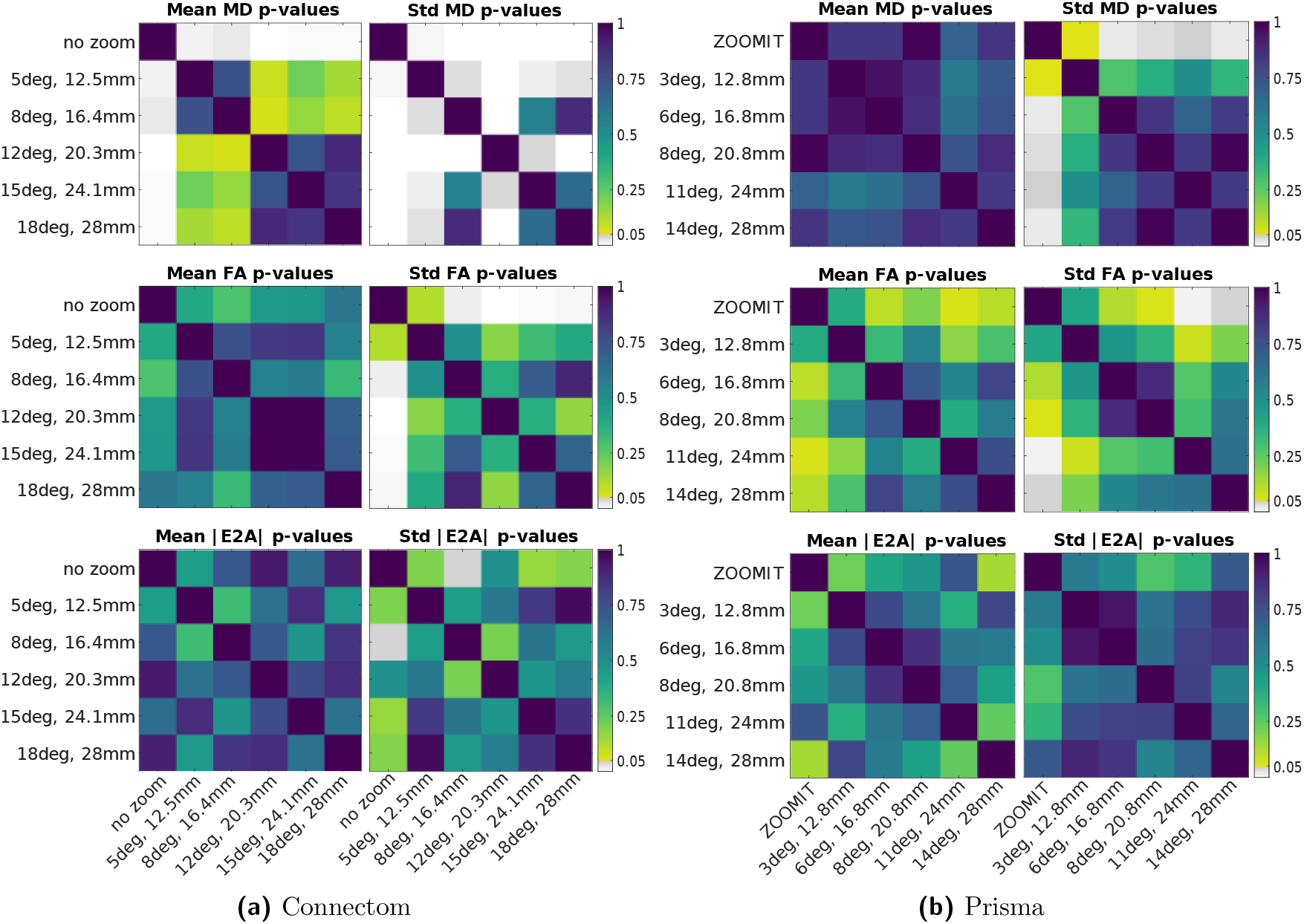
p-values comparing the average and standard deviation of diffusion measures in each boxplot from Figures 8 and 9 .

On Prisma, Figures 8 b and 10 b shows no statistically significant differences on mean DTI measures, although we note that FA seems slightly elevated for ZOOM compared to ZOOMit. Figures 9 b and 10 b show that the standard deviation over voxels shows a statistically significant decrease for ZOOMit compared to (most of) the ZOOM settings for MD and some ZOOM settings for FA. For |E2A|, although changes are visible in both examples Figures 6 and 7, we found no statistically significant differences overall for |E2A| on either scanner.

## 4 DISCUSSION

The ZOOM sequence provides a suitable method for acquiring reduced FOV diffusion weighted images of the human heart in vivo. Without ZOOM, one must increase the FOV for cardiac applications in the phase-encoding direction to avoid aliasing artifacts. The larger FOV results in a longer EPI readout increasing both susceptibility induced distortions and minimum achievable TE, and therefore adversely affects image quality and SNR. Using a limited FOV we were able to acquire cardiac diffusion weighted images with a short echo time of TE = 59ms on Connectome, and on Prisma it was possible to achieve TE = 67ms with ZOOM. Our results comparing ZOOM to ZOOMit on Prisma demonstrate that ZOOM can achieve image quality comparable to ZOOMit localization. Our results on Connectom (where ZOOMit is not available) show that ZOOM can be used to obtain DWI images with substantially lower artifact levels than acquisitions without ZOOM. Notably, on the Connectom system, ZOOM acquisitions yielded significantly lower average MD values and lower standard deviation over MD and FA, indicating reduced artifacts relative to scans without ZOOM. The lack of statistically significant differences between different ZOOM configurations supports the robustness of ZOOM to variations in ZOOM parameters over the region of parameter space identified using phantom scans. We selected ZOOM settings on the basis that the different settings would provide similar results to each other (due to being selected from an optimum ‘valley’ of ZOOM settings determined in phantom experiments before). Although we did not detect statistically significant differences in average diffusion measures between ZOOM settings, this should not be interpreted as meaning that these differences do not exist, but rather than our study is probably underpowered to detect these differences (any differences are obscured by inter-scan variability). While this makes it impossible for us to recommend a generalized, single particular ZOOM setting from this study, we emphasize that our hypothesis was that there are many suitable choices of ZOOM settings. The phantom experiments do not necessarily give the best settings for in vivo experiments. It is reasonable to assume that the optimal setting is subject dependent, and that choosing an optimal ZOOM setting for an individual may be guided by scout scans (just as placing the FOV and saturation bands may require scout scans). For an in vivo parameter search, on Prisma it is conceivable that ZOOMit could be use as a reference, and a similar parameter search performed as we have done with phantom scans, e.g., for a single diffusion weighting, obtain images for each ZOOM setting (and ZOOMit) and construct an RMSD surface in ZOOM parameter space. There may be subtle differences in slice profiles, SAR behavior, and excitation efficiency between ZOOM and ZOOMit, the consequences of which could be investigated with such an approach. For the Connectom scanner, there is no suitable reference scan to be used, making optimal parameter selection using in vivo scans more difficult. However, it is possible that an in vivo full-FOV acquisition could be used as a reference against ZOOM with a reduced-FOV, so long as a suitable image comparison was used that compensated for differences in echo times etc. ZOOM provides the additional advantage over ZOOMit due to the shorter RF excitation pulse duration, lowering the minimum achievable TE. On Prisma, it would have been possible to lower TE from 79ms with ZOOMit to 67ms with ZOOM, but we kept TE at 79ms in order to reduce confounding factors in our comparison between ZOOMit and ZOOM on Prisma. Assuming T2 *∼*46ms, ^25^ switching from ZOOMit to ZOOM could potential provide an SNR boost of 30%. For center-out readout techniques such as spiral, ^43,44^ our previous work withspiral readout on Prisma gives TE of 75ms with ZOOMit and 56ms with slice selective excitation (which would match TE for ZOOM), giving an estimated SNR increase of 50%. ^45^ For Connectom, we used TE = 59ms for ZOOM since we did not have ZOOMit to compare against. While the ultra-high gradients already allow for a lower TE, when combined with a spiral readout it is feasible that TE could approach T2 (at 3T) of 46ms in the heart.

## 5 CONCLUSION

We demonstrate that ZOOM is a viable alternative to 2D selective RF-pulses to achieve a reduced FOV acquisition in cDTI. This allowed us to achieve a shorter TE on a specialized scanner with ultra-strong gradient system for cDTI acquisitions. Sequences utilizing center-out readout techniques will particularly benefit from the echo time savings and, hence, SNR boost. Similarly, ZOOM will also be suitable for application of cDTI on MR systems with field strengths *≥*7T, which will require using the shortest possible TEs.

## 6 ACKNOWLEDGMENTS

For the purpose of open access, the author has applied a CC BY public copyright license to any Author Accepted Manuscript version arising from this submission. We thank Siemens Healthineers for the pulse sequence development environment.

## Author contributions

LM: conceptualization, sequence implementation, original draft; SC: experimental design, post-processing tools, data analysis, original draft; IW: data collection, data analysis, original draft; AD: conceptualization; IT: proof reading; ML: proof reading; ED: supervision, proof reading; FS: sequence implementation, proof reading DJ: supervision; MA: conceptualization, data collection, data analysis, original draft; JS: conceptualization, supervision, proof reading, funding

